# USP28 deletion and small molecule inhibition destabilises c-Myc and elicits regression of squamous cell lung carcinoma

**DOI:** 10.1101/2020.11.17.377705

**Authors:** E. Josue Ruiz, Adan Pinto-Fernandez, Andrew P. Turnbull, Linxiang Lan, Thomas M. Charlton, Hannah Claire Scott, Andreas Damianou, George Vere, Eva M. Riising, Clive Da Costa, Wojciech W. Krajewski, David Guerin, Jeffrey Kearns, Stephanos Ioannidis, Marie Katz, Crystal McKinnon, Jonathan C. O’Connell, Natalia Moncaut, Ian Rosewell, Emma Nye, Neil Jones, Claire Heride, Malte Gersch, Min Wu, Christopher J. Dinsmore, Tim R. Hammonds, Sunkyu Kim, David Komander, Sylvie Urbé, Michael J. Clague, Benedikt M. Kessler, Axel Behrens

**Affiliations:** Adult stem cell laboratory; The Francis Crick Institute, 1 Midland Road, London NW1 1AT, UK; Target Discovery Institute, Nuffield Department of Medicine, University of Oxford, Roosevelt Drive, Oxford OX3 7FZ, UK; CRUK Therapeutic Discovery Laboratories, The Francis Crick Institute, 1 Midland Road, London NW1 1AT, UK; FORMA Therapeutics, Arsenal Street, Watertown, Massachusetts 02472, USA; Genetic Manipulation Service, The Francis Crick Institute, 1 Midland Road, London NW1 1AT, UK; Max Planck Institute of Molecular Physiology, Otto-Hahn-Str 11, 44227 Dortmund, Germany; Ubiquitin Signalling Division, Walter and Eliza Hall Institute of Medical Research, Royal Parade, Parkville VIC 3052, and Dept of Medical Biology, University of Melbourne, VIC 3010, Australia; Cellular and Molecular Physiology, Institute of Translational Medicine, University of Liverpool, Crown Street, Liverpool L69 3BX, UK; Cancer Stem Cell Laboratory, Institute of Cancer Research, London, UK; Imperial College, Division of Cancer, Department of Surgery and Cancer, London, UK; Convergence Science Centre, Imperial College, London, SW7 2BU, UK; Constellation Pharmaceuticals, 215 First St, Cambridge, MA 02142, USA; Novartis Institutes for BioMedical Research, 250 Massachusetts Ave, Cambridge, MA 02139, USA; H3 Biomedicine, 300 Technology Square, Cambridge, MA 02139, USA; Valo Health, 399 Boylston St, Suite 505, Boston, MA 02116, USA; Disc Medicine, 150 Cambridgepark Drive Suite 103, Cambridge, MA 02135, USA; Kronos Bio, Inc., 301 Binney Street, 2nd Floor East, Cambridge, MA 02142, USA; Locki Therapeutics, London Bioscience Innovation Centre, 2 Royal College Street, London NW1 0NH, UK; Incyte, 1801 Augustine Cut-off, Wilmington, DE 19803, USA

## Abstract

Lung squamous cell carcinoma (LSCC) is a considerable global health burden, with an incidence of over 600,000 cases per year. Treatment options are limited, and patient 5-year survival rate is less than 5%. The ubiquitin specific protease 28 (USP28) has been implicated in tumorigenesis through its stabilization of the oncoprotein c-MYC. Here, we show that genetic inactivation of *Usp28* induced regression of established murine LSCC lung tumors. We developed a small molecule that inhibits USP28 activity in the low nanomole range. While displaying cross-reactivity against the closest homologue USP25, this inhibitor showed a high degree of selectivity over other deubiquitinases. USP28 inhibitor treatment resulted in a dramatic decrease in c-Myc proteins levels and consequently induced substantial regression of autochthonous murine LSCC tumors and human LSCC xenografts, thereby phenocopying the effect observed by genetic deletion. Thus, USP28 may represent a promising therapeutic target for the treatment of squamous cell lung carcinoma.

## Introduction

Lung cancer is the leading cause of cancer death worldwide. Based on histological criteria lung cancer can be subdivided into non-small cell lung cancer (NSCLC) and the rarer small cell lung cancer. The most common NSCLCs are lung adenocarcinoma (LADC) and squamous cell carcinoma (LSCC), with large cell carcinoma being less commonly observed. Progress has been made in the targeted treatment of LADC, largely due to the development of small-molecule inhibitors against EGFR, ALK, and ROS1 (Cardarella and Johnson, 2013). However, no targeted treatment options exist for LSCC patients (Hirsch et al., 2017; Novello et al., 2014). Consequently, despite having limited efficacy on LSCC patient survival, platinum-based chemotherapy remains the cornerstone of current LSCC treatment (Fennell et al., 2016; Isaka et al., 2017; Scagliotti et al., 2008). Therefore, there is an urgent need to identify novel druggable targets for LSCC treatment and to develop novel therapeutics.

The *FBXW7* protein product F-box/WD repeat-containing protein 7 (FBW7) is the substrate recognition component of an SCF-type ubiquitin ligase, which targets several well-known oncoproteins, including c-Myc, Notch, and c-Jun, for degradation (Davis et al., 2014). These oncoproteins accumulate in the absence of FBW7 function, and genetic analyses of human LSCC samples revealed common genomic alterations in *FBXW7* (Cancer Genome Atlas Research, 2012; Kan et al., 2010). In addition, FBW7 protein is undetectable by immunohistochemistry (IHC) in 69% of LSCC patient tumor samples (Ruiz et al., 2019). Genetically engineered mice harboring loss of *Fbxw7* concomitant with *KRasG12D* activation (KF mice) develop LSCC with 100% penetrance and short latency, as well as LADC (Ruiz et al., 2019). Thus, FBW7 is an important tumor suppressor in both human and murine lung cancer.

The deubiquitinase USP28 opposes FBW7-mediated ubiquitination of the oncoproteins c-Myc and c-Jun, thereby stabilizing these proteins (Popov et al., 2007). In a murine model of colorectal cancer, deleting *Usp28* reduced size of established tumors and increased lifespan (Diefenbacher et al., 2014). Therefore, targeting USP28 in order to destabilize its substrates represents an attractive strategy to inhibit the function of c-Myc and other oncogenic transcription factors that are not amenable to conventional inhibition by small molecules.

Here, we describe the characterisation of a novel USP28 inhibitory compound (USP28i) and the genetic as well as chemical validation of USP28 as a promising therapeutic target for LSCC tumors. Using an FRT/FLP and CRE/LOXP dual recombinase system (Schonhuber et al., 2014), we show that *Usp28* inactivation in established LSCC results in dramatic tumor regression. Importantly, USP28i treatment recapitulates LSCC regression in both mouse models and human LSCC xenografts. Absence or inhibition of USP28 resulted in a dramatic decrease in the protein levels of c-Myc, providing a potential mechanism of action for USP28i. Therefore, USP28 inhibition should be a strong candidate for clinical evaluation, particularly given the paucity of currently available therapy options for LSCC patients.

## Results

### USP28 is required to maintain protein levels of c-Myc, c-Jun and Δp63 in LSCC

To gain insights into the molecular differences between LADC and LSCC, we investigated the expression of MYC in these common NSCLCs subtypes. MYC was transcriptionally upregulated in human LSCC compared to healthy lung tissue or LADC tumors (**Figure 1A**). Quantitative polymerase chain reaction (qPCR) analysis on an independent set of primary human lung biopsy samples confirmed that MYC is highly expressed in LSCC tumors compared with normal lung tissue (**Figure 1B**). Moreover, immunohistochemistry (IHC) staining on primary lung tumors confirmed a significant abundance of c-Myc protein in LSCC samples (**Figure 1C, 1D**). Also Δp63 and c-Jun, critical factors in squamous cell identity and tumor maintenance, respectively, showed higher protein levels in LSCC compared to LADC tumors (**Figure 1C, 1D**). Individual downregulation of c-Myc, c-Jun and Δp63 by siRNA resulted in a significant reduction of cell growth in four independent human LSCC cell lines (**Figure 1E, S1A-C**).

**Figure 1.**
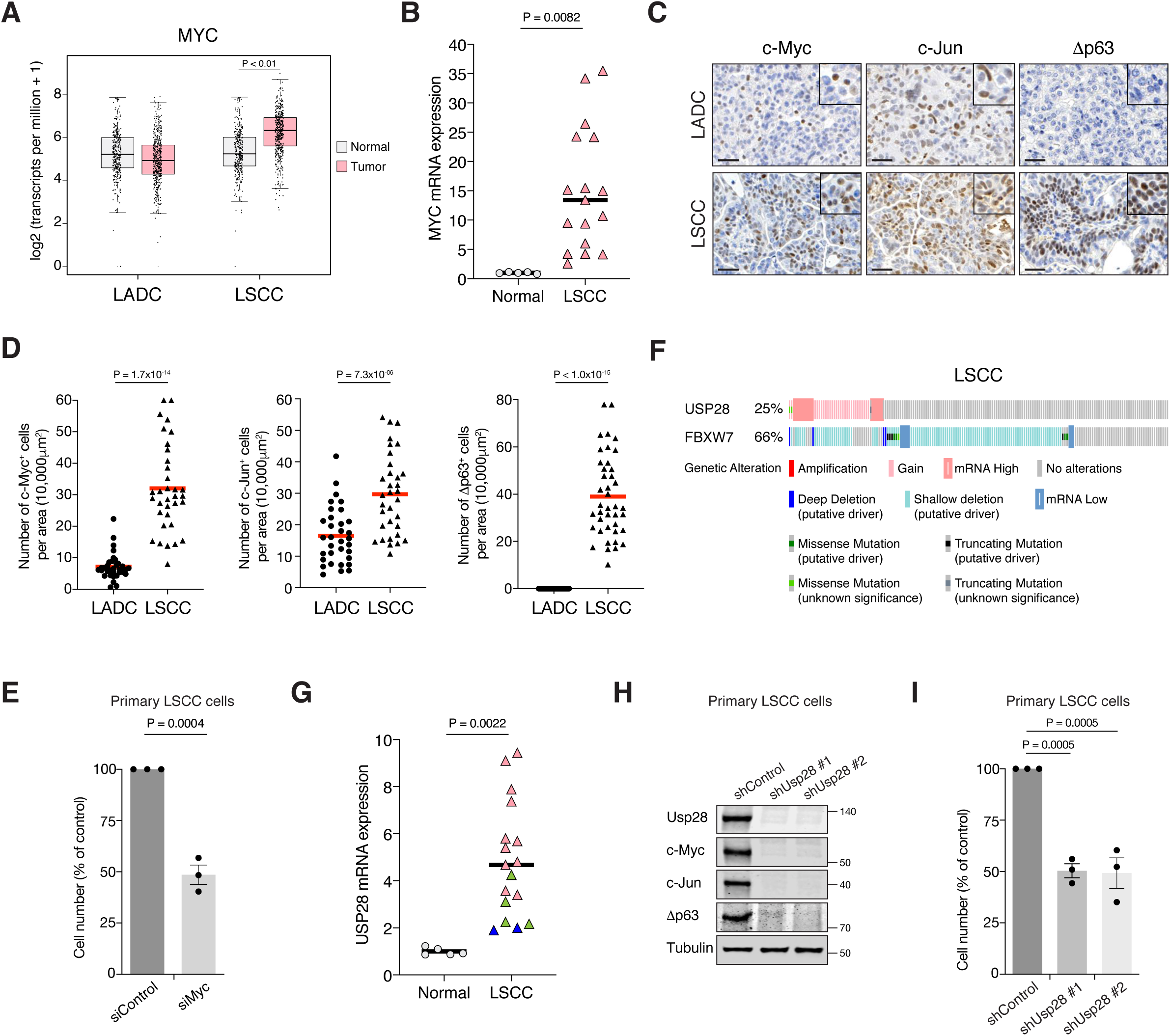
**MYC, JUN and Δp63 are highly expressed in LSCC tumors** A) Expression of *MYC* in human lung adenocarcinoma (LADC, n = 483), lung squamous cell carcinoma (LSCC, n = 486), and normal non-transformed tissue (normal LSCC = 338, normal LADC = 347). In box plots, the centre line reflects the median. Data from TCGA and GTEx were analyzed using GEPIA software. B) Relative mRNA expression of *MYC* in normal lung tissue (n = 5) and LSCC (n = 17) patient samples from the Cordoba Biobank measured by RT-PCR. The P value was calculated using the Student’s two-tailed t test. Plots indicate mean. C) Representative LADC and LSCC tumors stained with c-Myc, c-Jun and Δp63 antibodies. Scale bars, 30 μm. D) Quantification of c-Myc^+^ (LADC n = 33, LSCC n = 34), c-Jun^+^ (LADC n = 33, LSCC n = 33) and Δp63^+^ cells (LADC n = 41, LSCC n = 41) in LADC and LSCC tumors. Plots indicate mean. Student’s two-tailed t test was used to calculate P values. E) Graph showing the difference in cell proliferation between control and MYC-depleted KF LSCC cells (n = 3). Graph indicates mean ± S.E.M.. Student’s two-tailed t test was used to calculate P values. F) Genetic alterations in *USP28* and *FBXW7* genes in human LSCC. Each column represents a tumor sample (n = 178). Data from TCGA were analyzed using cBioportal software. G) Relative mRNA expression of *USP28* in normal lung tissue (n = 5) and LSCC (n = 17) patient samples from the Cordoba Biobank measured by RT-PCR. The P value was calculated using the Student’s two-tailed t test. Plots indicate mean. See also Supplementary Figure S2B. H) shRNA-mediated knockdown of Usp28 decreases c-Myc, c-Jun and Δp63 protein levels in primary KF LSCC cells. I) Graph showing the difference in cell proliferation between control and Usp28-depleted KF LSCC cells (n = 3). Graph indicates mean ± S.E.M.. One-way ANOVA with Dunnett’s multiple comparisons test was used to calculate P values.

As c-Myc, c-Jun and Δp63 protein levels are controlled by the deubiquitinase USP28 (Popov et al., 2007; Prieto-Garcia et al., 2020), we analysed its expression in publicly available datasets (The Cancer Genome Atlas). We observed that 25% of human LSCC cases show gain-of-function alterations in *USP28* (**Figure 1F**). In addition, a positive correlation between *USP28* copy-number and mRNA expression was found in the same datasets (**Figure S2A**). Interestingly, qPCR and IHC analysis on human LSCC samples revealed that low *USP28* mRNA levels correlated with low USP28 protein levels and likewise, high/moderate mRNA levels also correlated with high USP28 protein levels (**Figure 1G, S2B**). Since USP28 is involved in Δp63, c-Jun and c-Myc stabilization and higher expression of USP28 is associated with a significantly shorter survival time (Prieto-Garcia et al., 2020), we targeted its expression. Usp28 downregulation by shRNA resulted in a significant reduction in c-Myc, c-Jun and Δp63 protein levels in LSCC primary tumor cells and reduced LSCC cell growth (**Figure 1H, 1I**). Thus, targeting USP28 in order to destabilize its substrates represents a rational strategy to target tumor cells that rely on oncogenic transcription factors that are currently not druggable by small molecules.

### Generation of a pre-clinical dual recombinase lung cancer mouse model

Recently, Usp28 was shown to be required for the initiation of lung tumors in the Rosa26-Cas9 sgRNA Kras^G12D^; Tp53; Lkb1 model (Prieto-Garcia et al., 2020). However, a meaningful pre-clinical model requires targeting the therapeutic candidate gene in existing growing lung tumors. Thus, to assess the function of *Usp28* in established tumors, we developed a new genetically engineered mouse (GEM) model to temporally and spatially separate tumor development from target deletion by using two independent recombinases: Flp and Cre^ERT^. In this model, LSCC and LADC formation is initiated by KRas^G12D^ activation and *Fbxw7* deletion using Flp recombinase, and the Cre/loxP system can then be used for inactivation of Usp28^flox/flox^ in established tumors. To allow conditional FRT/Flp-mediated inactivation of *Fbxw7* function, we inserted two FRT sites flanking exon 5 of the endogenous *Fbxw7* gene in mice to generate a Fbxw7^FRT/FRT^ allele that can be deleted by Flp recombinase (**Figure S3A, S3B**). Expression of Flp recombinase resulted in the deletion of *Fbxw7* exon 5, which could be detected by PCR (**Figure S3B**). The resulting strain, Fbxw7^FRT/FRT^, was crossed to FRT-STOP-FRT (FSF)-KRas^G12D^ mice to generate FSF-KRas^G12D^; Fbxw7^FRT/FRT^ (KF-Flp model).

### USP28 is an effective therapeutic target for LSCC, but not KRas^G12D^; Trp53 mutant LADC tumors

The KF-Flp strain described above was crossed with ROSA26-FSF-Cre^ERT^; Usp28^flox/flox^ mice to generate the KFCU model (**Figure 2A**). KFCU tumor development was monitored by CT scans. At ten-to-eleven weeks post-infection with Flp recombinase-expressing recombinant adenoviruses, animals displayed lesions in their lungs. At this time point, we confirmed by histology that KFCU mice develop both LADC and LSCC tumors (**Figure S3C**). As expected (Ruiz et al., 2019), KFCU LADC lesions occurred in alveolar tissue and were positive for Sftpc and TTF1. KFCU LSCC tumors occurred mainly in bronchi (rarely manifesting in the alveolar compartment) and expressed CK5 and Δp63. Next, animals displaying lung tumors were exposed to tamoxifen to activate the Cre^ERT^ protein and delete the conditional *Usp28* floxed alleles (**Figure 2A, S3D**). Although the loss of Usp28 expression decreased LADC tumor size, it did not reduce the number of LADC tumors (**Figure 2B-D**). In contrast, histological examination of KFCU mice revealed a clear reduction in the numbers of LSCC lesions in *Usp28*-deleted lungs (**Figure 2F, S3D**). As well as a significant reduction in tumor number, the few CK5-positive LSCC lesions that remained were substantially smaller than control tumors (**Figure 2G**). Measurement of the size of 429 individual KFCU LSCC tumors (326 vehicle-treated and 103 tamoxifen-treated) showed an average size of 11.4×10^4^ μm^2^ in the vehicle arm versus 4.6×10^4^ μm^2^ in the tamoxifen arm (**Figure 2G).** Thus, *Usp28* inactivation significantly reduces both the number and the size of LSCC tumors.

**Figure 2.**
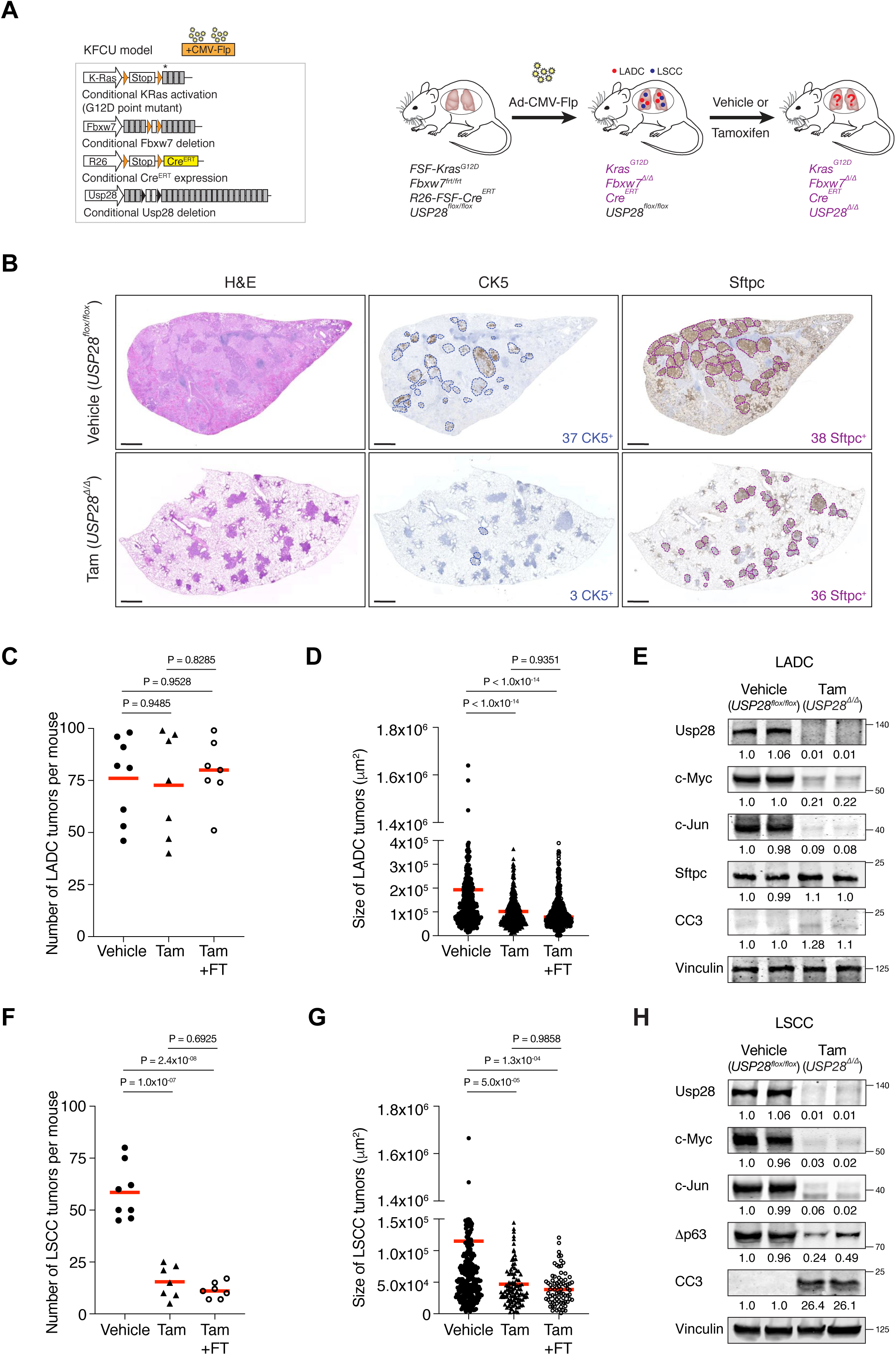
**Usp28 is an effective therapeutic target for LSCC tumors** A) Schematic representation of the KFCU (FSF-<u>K</u>ras^G12D^; <u>F</u>bxw7^FRT/FRT^; ROSA26-FSF-<u>C</u>re^ERT^; <u>U</u>sp28^flox/flox^) model and experimental approach used to deplete conditional Usp28 alleles in established lung tumors. B) Lung histology of animals treated as in A, showing both LSCC (CK5^+^) and LADC (Sftpc^+^) tumors in mice receiving vehicle but few LSCC lesions in mice receiving tamoxifen. Scale bars, 1000 μm. C) Quantification of LADC tumors in vehicle-, tamoxifen- and tamoxifen+FT206 treated KFCU mice. Plots indicate mean. One-way ANOVA with Tukey’s multiple ‘comparisons test was used to calculate P values (n = 8 vehicle, n = 7 tamoxifen, n = 7 tamoxifen + FT206). D) Quantification of LADC tumor size in vehicle-, tamoxifen- and tamoxifen+FT206 treated KFCU mice. Plots indicate mean. One-way ANOVA with Tukey’s multiple ‘comparisons test was used to calculate P values (n = 466 vehicle, n = 434 tamoxifen, n = 503 tamoxifen + FT206). E) Immunoblot analysis of LADC tumors probed for Usp28, c-Myc, c-Jun, Sftpc, cleaved caspase-3 (CC3). Vinculin is shown as loading control. F) Quantification of LSCC tumors in vehicle-, tamoxifen- and tamoxifen+FT206 treated KFCU mice. Plots indicate mean. One-way ANOVA with Tukey’s multiple ‘comparisons test was used to calculate P values (n = 8 vehicle, n = 7 tamoxifen, n = 7 tamoxifen + FT206). G) Quantification of LSCC tumor size in vehicle-, tamoxifen- and tamoxifen+FT206 treated KFCU mice. Plots indicate mean. One-way ANOVA with Tukey’s multiple ‘comparisons test was used to calculate P values (n = 326 vehicle, n = 103 tamoxifen, n = 79 tamoxifen + FT206). H) *Usp28* deletion induces apoptotic cell death (cleaved caspase-3, CC3) and decreases c-Myc, c-Jun and Δp63 protein levels in LSCC lesions.

To get insights into LSCC tumor regression, we focused on Usp28 substrates. Immunoblotting analysis revealed that *Usp28* deletion resulted in apoptotic cell death (cleaved caspase-3; CC3). Δp63 protein levels were reduced, but c-Jun and c-Myc protein became undetectable (**Figure 2H, S3E**). *Usp28* deletion also decreased c-Jun and c-Myc levels in KFCU LADC lesions, although the reduction in c-Myc protein levels were significantly less pronounced than observed in LSCC (**Figure 2E**). Strikingly, elimination of *Usp28* has little effect, if any, on apoptotic cell death, as determined by its inability to induce cleaved caspase-3 in LADC lesions. Thus, these data suggest that Usp28 and its substrates are required for the maintenance of LSCC tumors.

To further investigate the role of Usp28 in LADC, we studied the consequences of *Usp28* deletion in a second LADC genetic model. We used Flp-inducible oncogenic K-Ras activation combined with p53 deletion (FSF-KRas^G12D^ and Trp53^FRT/FRT^ or KP-Flp model) (Schonhuber et al., 2014). The KP-Flp mice were crossed to a conditional Usp28^flox/flox^ strain together with an inducible Cre^ERT^ recombinase knocked in at the ROSA26 locus and an mT/mG reporter allele (KPCU mice; **Figure 3A**). After intratracheal adeno-CMV-Flp virus instillation, *Usp28* was deleted in KPCU animals displaying lung tumors by CT (**Figure 3A**). Loss of Usp28 expression in this second LADC model also did not result in a reduction of LADC tumor number and size (**Figure 3C, 3D**). Successful Cre^ERT^ recombination was verified using lineage tracing (GFP staining) and deletion of Usp28^flox/flox^ alleles was further confirmed by BaseScope assays (**Figure 3B, 3E**). Therefore, also these data argue against an important role for Usp28 in LADC tumors.

**Figure 3.**
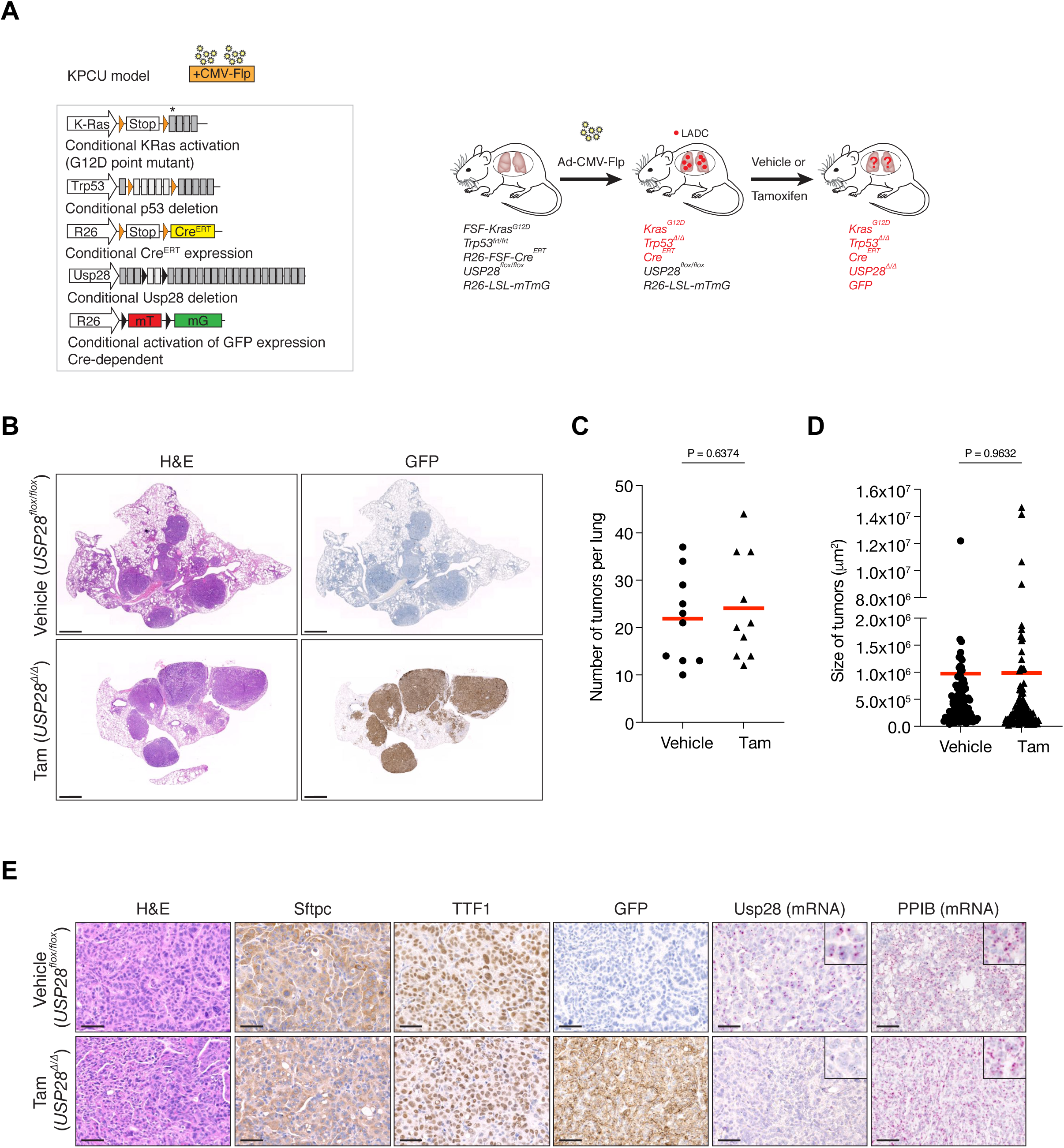
**Usp28 is not a therapeutic target for advanced KRas^G12D^; Trp53 mutant tumors** A) Schematic representation of the KPCU (FSF-<u>K</u>Ras^G12D^; <u>p</u>53^FRT/FRT^; ROSA26-FSF-<u>C</u>re^ERT^; <u>U</u>sp28^flox/flox^; ROSA26-LSL-mTmG) model and experimental approach used. At 10-weeks post-infection, KPCU mice were treated with vehicle or tamoxifen. B) Representative images of H&E (left) and GFP (right) stains from mice of the indicated treatments. Scale bar, 1000 µm. C) Quantification of mouse LADC tumors in the KPCU model. Plots indicate mean. Student’s two-tailed t test was used to calculate P values (n = 10 vehicle, n = 10 tamoxifen). D) Quantification of LADC tumor size in vehicle- and tamoxifen-treated KPCU mice. Plots indicate mean. Student’s two-tailed t test was used to calculate P values (n = 110 vehicle, n = 130 tamoxifen). E) Representative images illustrating histological analysis of lung lesions in KPCU mice, treated with vehicle or tamoxifen. H&E, Sftpc, TTF1, GFP immunohistochemistry staining and in situ hybridization of USP28 and PPIB mRNA expression. Scale bars, 50 µm.

### Generation of a new USP28 inhibitor: selectivity and cellular target engagement

The finding that Usp28 plays a key role in LSCC tumor maintenance prompted us to identify small molecule inhibitors against this deubiquitinase. A small molecule discovery campaign based on the ubiquitin-rhodamine cleavable assay (Turnbull et al., 2017) yielded a panel of compounds sharing a thienopyridine carboxamide chemical scaffold with inhibitory selectivity for USP28 and USP25 (Guerin, 2017; Guerin et al., 2020; Zablocki et al., 2019). The compound FT206 (**Figure 4A**) represents a different chemical class from the benzylic amino ethanol-based inhibitors described previously (Wrigley et al., 2017). Quantitative structure-activity relationship (SAR) was used to develop compound derivative FT206 that was most optimal in terms of drug metabolism and pharmacokinetic properties (DMPK) while preserving potency and selectivity towards USP28/25 (Zablocki et al., 2019). To confirm FT206 cellular target engagement, we used a Ub activity-based probe assay (ABP) (Altun et al., 2011; Clancy et al., 2021; Panyain et al., 2020; Turnbull et al., 2017). ABPs can assess DUB enzyme activity in a cellular context. DUB inhibition leads to displacement of the ABP probe, resulting in a molecular weight shift measurable by SDS-PAGE and immunoblotting against USP28/25. Using this approach, we found that the compound FT206 interferes with USP28/25 probe labelling (USP-ABP versus USP) in LSCC H520 cell extracts (EC_50_ ∼300-1000nM, **Figure 4B**) and intact cells (EC_50_ ∼1-3μM, **Figure 4C**). In contrast to FT206, AZ1, a different USP28 inhibitor (Wrigley et al., 2017), based on a benzylic amino ethanol scaffold, appeared to exert lower potency towards USP28 (EC_50_ >30μM) and selectivity for USP25 (EC_50_ ∼10-30μM) (**Figure S4A**). To address compound selectivity more widely, we combined the ABP assay with quantitative mass spectrometry (ABPP) to allow the analysis of the cellular active DUBome (Benns et al., 2021; Jones et al., 2021; Pinto-Fernandez et al., 2019). When performing such assay in human LSCC cells, we were able to profile 28 endogenous DUBs, revealing a remarkable USP28/25 selectivity for FT206 in a dose-dependent manner (**Figure 4D**).

**Figure 4.**
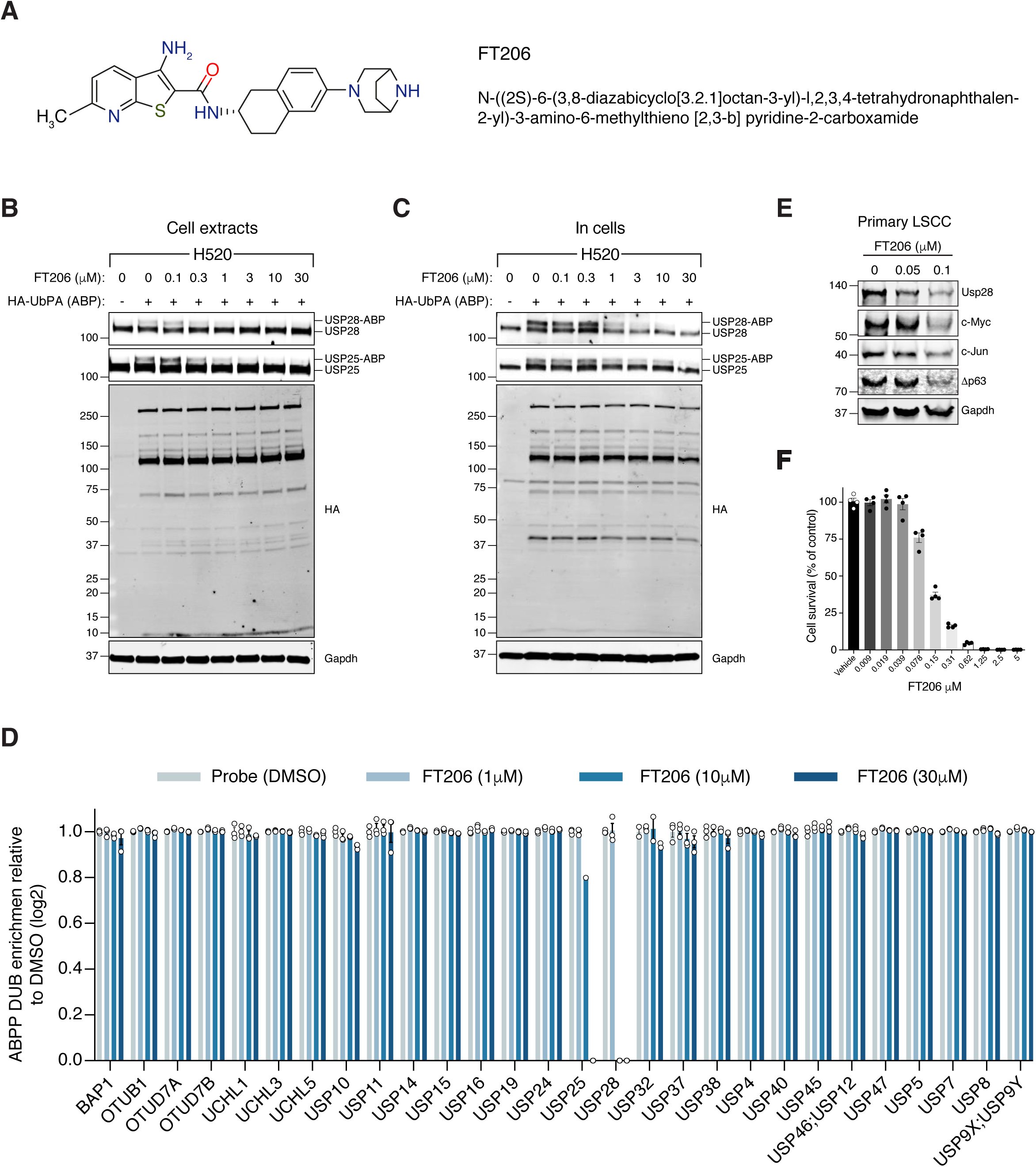
**USP28 inhibitor selectivity and cellular target engagement** A) Structure of small molecular inhibitor FT3951206/CRT0511973 (FT206). B) Cellular DUB profiling in NCI-H520 LSCC cell extracts incubated with the indicated concentrations of FT206 prior to labelling with HA-UbPA, SDS-PAGE and analysis by Western blotting. Inhibitor potency was reflected by competition with USP28/25-ABP adduct formation. C) Cellular DUB profiling in NCI-H520 LSCC cells incubated with the indicated concentrations of FT206, lysed extracts labelled with HA-UbPA and analysed as in B. D) Activity-based Probe Profiling (ABPP) demonstrating the cellular DUB selectivity profile of cpd FT206 by quantitative mass spectrometry analysis at different inhibitor concentrations. Graph indicates mean ± S.E.M.. E) Usp28 inhibition using FT206 (50nM and 100nM) reduces c-Myc, c-Jun and Δp63 protein levels in primary KF LSCC cells. F) Usp28 inhibition using FT206 decreases cell proliferation in KF LSCC cells (n = 4). Graph indicates mean ± S.E.M..

To further evaluate the efficacy of FT206 in targeting USP28, we tested its ability to modulate the ubiquitination status of endogenous USP28 substrates. The ubiquitination levels of c-Myc and c-Jun increased upon FT206 and MG132 co-treatment (**Figure 4SB**), confirming that FT206 blocks USP28-mediated deubiquitination of its substrates. The ubiquitination level of USP28 also increased upon FT206 treatment (**Figure 4SB**), which is consistent with previous observations where the enzymatic activity of DUBs can function to enhance their own stability (de Bie and Ciechanover, 2011). Consequently, treatment of LSCC tumor cells with FT206 resulted in reduced c-Myc, c-Jun, Δp63 and Usp28 protein levels, which were restored upon addition of MG132 (**Figure 4E, S4C**).

Finally, FT206 treatment impaired LSCC cell growth (**Figure 4F**). However, in a USP28-depleted background, FT206 neither affected cell growth nor reduced c-Myc protein levels (**Figure S4D**). Thus, this data suggests that the effects of FT206 are mediated by USP28.

### Pharmacological inhibition of USP28 is well tolerated in mice and induced LSCC tumor regression

We next evaluated the therapeutic potential of the USP28 inhibitor FT206 using the LSL-KRas^G12D^; Fbxw7^flox/flox^ model (KF mice), which develop both LADC and LSCC tumor types (Ruiz et al., 2019). Nine weeks after adeno-CMV-Cre virus infection, when mice had developed lung tumors, we started treatment with USP28 inhibitor at 75 mg/kg, 3 times a week for 5 weeks (**Figure 5A**). FT206 administration had no noticeable adverse effects and treated mice maintained normal body weight (**Figure S5A, S5B**). Consistent with the effects observed by genetic *Usp28* inactivation (**Figure 2C**), the number of KF LADC lesions was not affected by Usp28 inhibition via FT206 treatment (**Figure 5B, 5C, 5D**). By contrast, we found that FT206 effectively reduced LSCC tumor number by 68% (31 to 10 LSCC tumors, **Figure 5B, 5E**).

**Figure 5.**
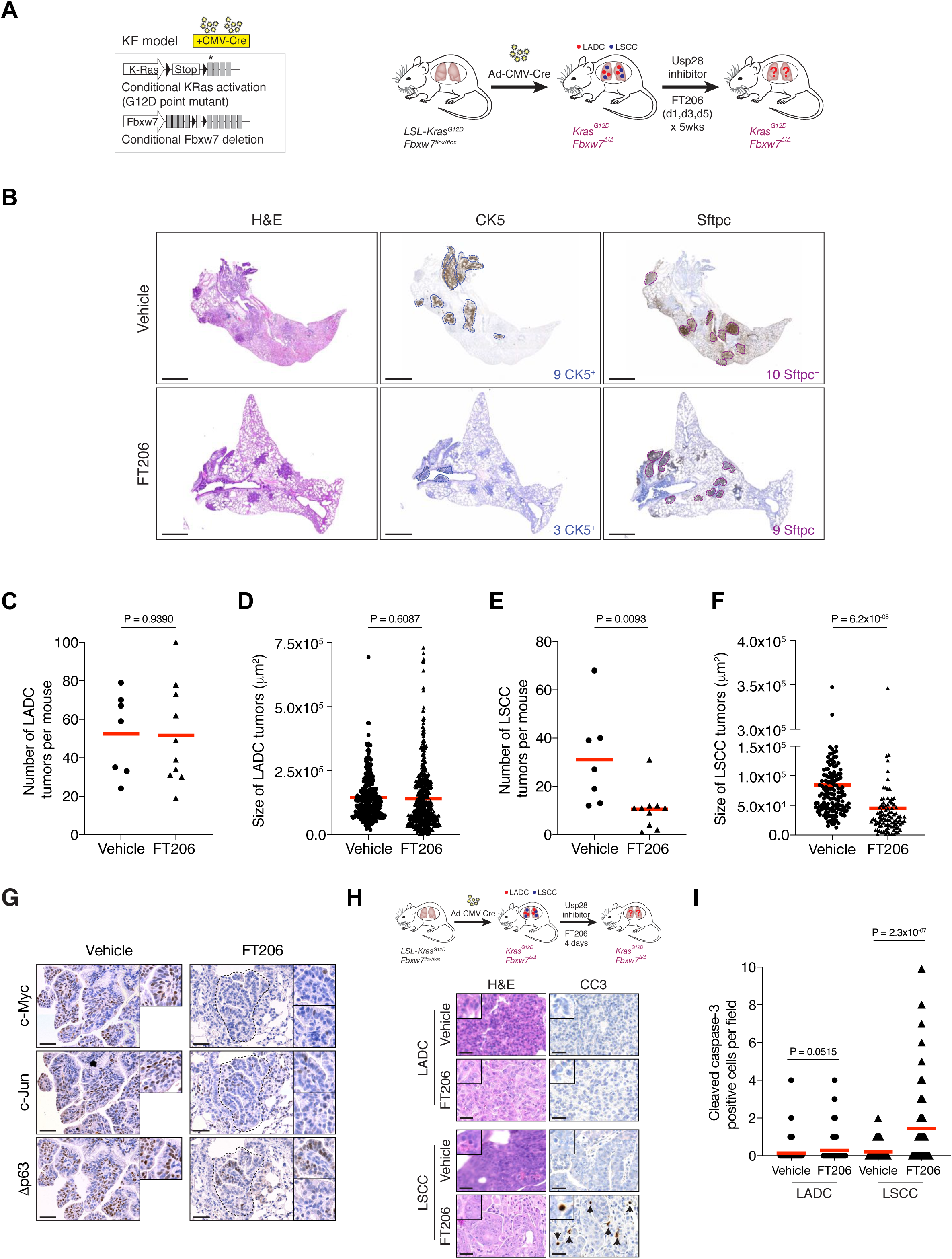
**Pharmacologic USP28 inhibition reduces c-Myc, c-Jun and Δp63 protein levels in mouse LSCC tumors, and induces tumor cell death** A) Scheme depicting experimental design for in vivo test of FT206 (75mg/kg), 3 times a week for 5 weeks. B) Lung histology of animals treated as in A, showing both LSCC (CK5^+^) and LADC (Sftpc^+^) tumors in LSL-KRas^G12D^; Fbxw7^f/f^ (KF) mice receiving vehicle but few LSCC lesions in mice receiving FT206. Scale bars, 1000 μm. C) Quantification of LADC tumors per animal in vehicle- and FT206-treated KF mice. Plots indicate mean. P values calculated using Student’s two-tailed t test (n = 7 vehicle, n = 10 FT206). D) Quantification of LADC tumor size in vehicle- and FT206-treated KF mice. Plots indicate mean. Student’s two-tailed t test was used to calculate P values (n = 304 vehicle, n = 481 FT206). E) Quantification of LSCC tumors per animal in vehicle- and FT206-treated KF mice. Plots indicate mean. P values calculated using Student’s two-tailed t test (n = 7 vehicle, n = 10 FT206). F) Quantification of LSCC tumor size in vehicle- and FT206-treated KF mice. Plots indicate mean. Student’s two-tailed t test was used to calculate P values (n = 156 vehicle, n = 96 FT206). G) LSCC tumors stained with c-Myc, c-Jun and Δp63 antibodies. KF animals treated with vehicle (left panel) or FT206 (right panel). Inserts showing c-Myc^+^, c-Jun^+^, Δp63^+^ LSCC tumors in mice receiving vehicle (left panel) but partial positive or negative LSCC lesions in mice receiving FT206 (right panel). Scale bars, 50 μm. H) Scheme depicting experimental design for in vivo test of FT206 (75 mg/kg) for 4 days consecutively (upper panel). Cleaved caspase-3 (CC3) stain shows apoptotic cells (bottom panel). Scale bars, 50 μm. I) Quantification of cleaved caspase-3 (CC3)-positive cells per field (20x) in LADC (n = 114 vehicle, 203 FT206) and LSCC (n = 94 vehicle, 167 FT206) tumors from KF mice treated as in H. Plots indicate mean. Student’s two-tailed t test was used to calculate P values. See also Supplementary Figure S5.

Moreover, measurement of 252 individual KF LSCC mutant tumors (156 vehicle-treated and 96 FT206-treated lesions) showed a significant reduction of over 45% in tumor size upon FT206 treatment: an average of 8.5×10^4^ μm^2^ in the vehicle arm versus 4.5×10^4^ μm^2^ in the FT206 cohort (**Figure 5F**). Thus, Usp28 inhibition by FT206 leads to a dramatic reduction in the numbers of advanced LSCC tumors, and the small number of remaining LSCC lesions are significantly reduced in size, resulting in a reduction of total LSCC burden of over 85% by single agent treatment.

In line with the effects found by genetic *Usp28* deletion, treatment of KF mice with FT206 also resulted in reduced Δp63, c-Jun and c-Myc protein levels (**Figure 5G**). Consequently, FT206 treatment led to a substantial increase in the number of cleaved caspase-3-positive cells in LSCC while LADC cells were not significantly affected, indicating that Usp28 inhibition causes apoptotic cell death of LSCC tumor cells (**Figure 5H, 5I**).

Finally, to further confirm the specificity of FT206, KFCU mice pre-exposed to tamoxifen to delete the conditional *Usp28* floxed alleles were further treated with the USP28 inhibitor FT206. In this setting, Usp28 inhibition did not result in a further reduction of LADC and LSCC lesions (**Figure 2C, 2F**), suggesting that FT206 targets specifically Usp28.

### USP28 inhibition causes dramatic regression of human LSCC xenograft tumors

To determine whether the promise of USP28 as a target in mouse lung cancer models can be translated to a human scenario, we established human xenograft tumor models. siRNA-mediated *USP28* depletion, and USP28 inhibitor treatment, considerably reduced protein levels of Δp63, c-Jun and c-Myc and impaired growth in human LSCC tumor cells (**Figure 6A-C, S6A**). In contrast, FT206 treatment had marginal effects on c-Myc and c-Jun protein levels in human LADC cells (**Figure S6B**). Crucially, FT206 led to a remarkable growth impairment of xenografts derived from three independent human LSCC cell lines (**Figure 6D-I**), which was accompanied with a strong reduction of c-Myc protein levels (**Figure 6J-L**). In summary, these data suggest that USP28 pharmacological intervention is a promising therapeutic option for human LSCC patients.

**Figure 6.**
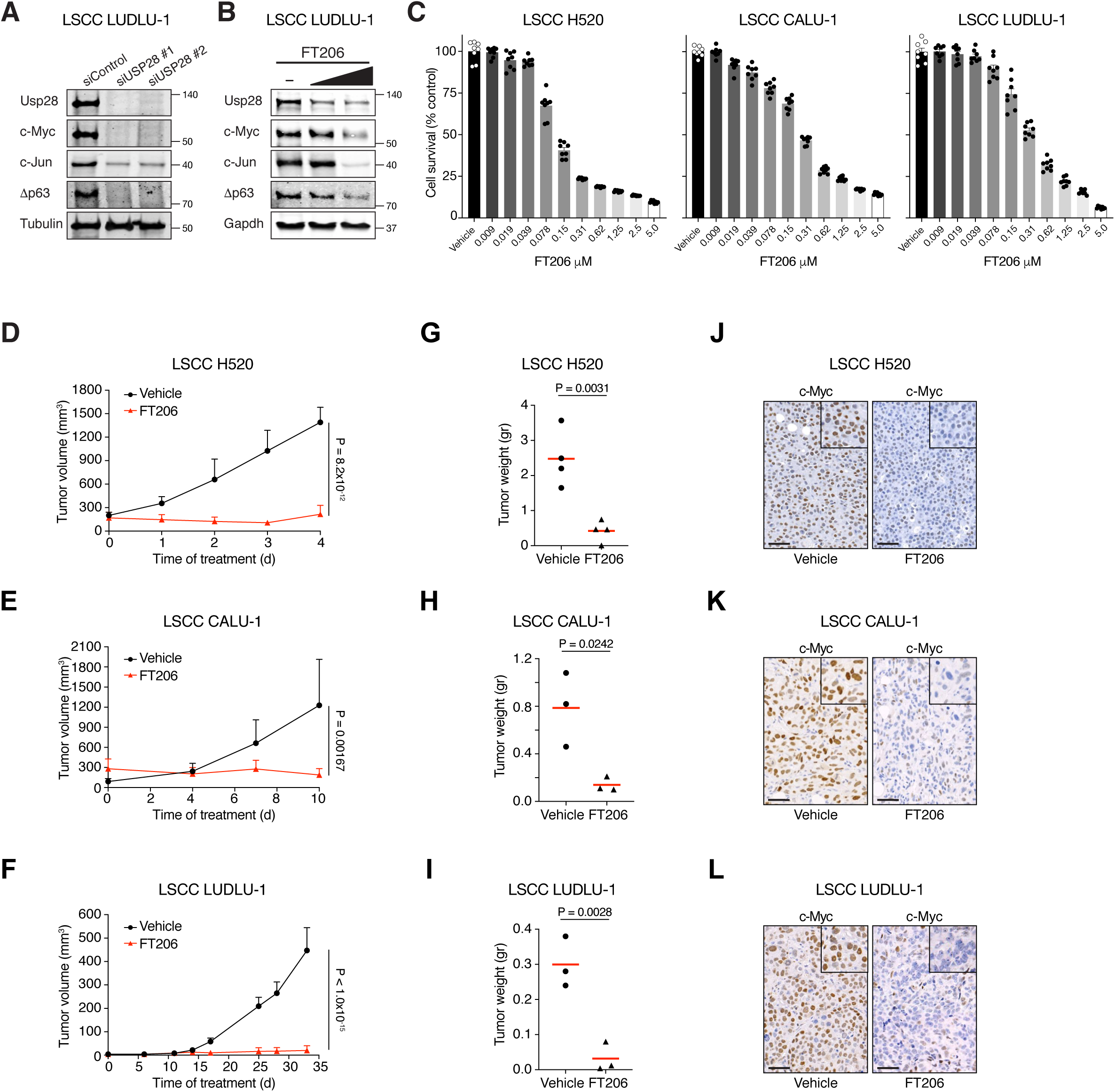
**Pharmacological inhibition of USP28 prevents human LSCC tumor progression and reduces c-Myc protein levels in xenograft models** A) siRNA-mediated knockdown of USP28 decreases c-Myc, c-Jun and Δp63 protein levels in human LUDLU-1 LSCC cells. B) USP28 inhibition using FT206 (0.2 and 0.4 μM) reduces c-Myc, c-Jun and Δp63 protein levels in human LUDLU-1 LSCC cells. C) USP28 inhibition using FT206 decreases cell proliferation in human LSCC (NCI-H520, CALU-1 and LUDLU-1) cell lines (n = 8). Graphs indicate mean ± S.E.M.. D, E, F) In vivo tumor graft growth curves of human LSCC (NCI-H520, CALU-1 and LUDLU-1) cell lines subcutaneously injected in flanks of immunocompromised mice. Animals with palpable tumors were treated with vehicle or FT206 (75mg/kg) via oral gavage. Plots indicate mean ± SD of the tumor volumes. P values calculated from two-way ANOVA with Bonferroni’s multiple comparisons test (NCI-H520 n = 4 vehicle and 4 FT206; CALU-1 n = 3 vehicle and 3 FT206; LUDLU-1 n = 3 vehicle and 3 FT206). G, H, I) Mice treated as in D, E and F, respectively. Plots showing the weight of xenograft tumors at the end point. Student’s two-tailed t test was used to calculate P values (NCI-H520 n = 4 vehicle and 4 FT206; CALU-1 n = 3 vehicle and 3 FT206; LUDLU-1 n = 3 vehicle and 3 FT206). J, K, L) c-Myc immunohistochemistry stainings of NCI-H520, CALU-1 and LUDLU-1 xenografts in mice treated as in D, E and F, respectively. Scale bars, 50 μm.

## Discussion

Unlike for LADC, there are few approved targeted therapies against LSCC. Consequently, despite its limited effectiveness on disease progression and prognosis, patients with LSCC receive the same conventional platinum-based chemotherapy today as they would have received two decades ago (Fennell et al., 2016; Gandara et al., 2015; Isaka et al., 2017; Liao et al., 2012; Scagliotti et al., 2008).

c-MYC is a transcription factor that orchestrates a potent pro-cancer programme across multiple cellular pathways. As c-MYC is often overexpressed in late-stage cancer, targeting it for degradation is an attractive strategy in many settings. The term ‘undruggable’ was coined to describe proteins that could not be targeted pharmacologically. Many desirable targets in cancer fall into this category, including the c-MYC oncoprotein, and pharmacologically targeting these intractable proteins is a key challenge in cancer research.

The deubiquitylase family of enzymes have emerged as attractive drug targets, that can offer a means to destabilize client proteins that might otherwise be undruggable (Schauer et al., 2019). The deubiquitinase USP28 was known to remove FBW7- mediated ubiquitination of, and thereby stabilise, the oncoprotein c-MYC (Popov et al., 2007). Importantly, mice lacking *Usp28* are healthy (Knobel et al., 2014), suggesting that Usp28 is dispensable for normal physiology and homeostasis.

In the current study we identified a requirement for USP28 for the maintenance of murine and human LSCC tumors. In agreement with the absence of major phenotypes in the *Usp28* knock out mice, USP28 inhibitor treatment was well tolerated by the experimental animals, while having a dramatic effect on LSCC regression. USP28 small molecule inhibition phenocopies the effects of *Usp28* deletion in LSCC regression, consistent with on-target activity. However, we cannot exclude that the inhibition of USP25 and possibly additional off-targets effects may contribute to the observed phenotype. Inhibitor treated mice kept a normal body weight, indicating no global adverse effects (**Figure S5A**).

While USP28 inhibition resulted in profoundly reduced LSCC growth, the effect on LADC was modest. TP63, c-Jun and c-Myc protein levels are increased in LSCC compared to LADC (**Figure 1C, 1D**). This could indicate a greater dependence of LSCC on these oncoproteins, which consequently may result in increased sensitivity to USP28 inhibition. We previously found that *Usp28* deficiency corrected the accumulation of SCF (Fbw7) substrate proteins, including c-Jun and c-Myc, in *Fbw7*- mutant cells (Diefenbacher et al., 2015). The frequent downregulation of *FBXW7* in human LSCC (Ruiz et al., 2019) (**Figure S2B**) may underlie the increased accumulation of SCF(Fbw7) substrate proteins like c-Myc, c-Jun and p63 in LSCC, and thereby cause LSCC tumors to be increasingly dependent on USP28 function. Indeed, our study suggest that those 3 oncoproteins are all relevant targets of USP28 in LSCC (**Figure 2H**). In contrast, Prieto-Garcia et al. saw no difference in c-Jun and c-Myc protein levels, and suggested a different mechanism of action. Of note, our and the Prieto-Garcia et al. studies used different dual specificity inhibitors of USP28/25 that have distinct properties. FT206, the compound used in this study, preferentially inhibits USP28 compared to USP25, whereas AZ1, the compound used by Prieto-Garcia et al. showed a pronounced activity towards USP25. In addition, FT206 inhibits USP28 in the nano-molar range, while Prieto-Garcia et al. typically used AZ1 at 10-30μM, possibly because higher compound concentrations are required for therapeutic inhibition of USP28. Therefore differences in the selectivity and potency of the compounds used may explain some of the differences observed.

Interestingly, all human LSCC cell lines used in the xenograft experiment (**Figure 6**), each of which responded well to USP28 inhibition, do not show neither gain- or loss- of-function mutations in *USP28* nor *FBXW7*, respectively. Thus, these data support the notion that LSCC tumor cells respond to USP28 inhibition, regardless of *USP28/FBXW7* mutation status, which suggest that USP28 inhibition might be a therapeutic option for many LSCC patients.

In summary, our studies demonstrate that USP28 is a key mediator of LSCC maintenance and progression and hence USP28 represents an exciting therapeutic target. Therefore, USP28 inhibition should be considered as a potential therapy for human lung squamous cell carcinoma.

## Methods and Materials

### Mice

The LSL-KRas^G12D^ (Jackson et al., 2001), Fbxw7^flox/flox^ (Jandke et al., 2011), Usp28^flox/flox^ (Diefenbacher et al., 2014), FSF-KRas^G12D^ (Schonhuber et al., 2014), Trp53^FRT/FRT^ (Schonhuber et al., 2014), ROSA26-FSF-Cre^ERT^ (Schonhuber et al., 2014), ROSA26-LSL-mTmG (Muzumdar et al., 2007) strains have been previously described. Immunocompromised NSG mice were maintained in-house. All animal experiments were approved by the Francis Crick Institute Animal Ethics Committee and conformed to UK Home Office regulations under the Animals (Scientific Procedures) Act 1986 including Amendment Regulations 2012. All strains were genotyped by Transnetyx. Each group contained at least 3 mice, which generates enough power to pick up statistically significant differences between treatments, as determined from previous experience (Ruiz et al., 2019). Mice were assigned to random groups before treatment.

### Generation of Fbxw7^FRT/FRT^ Mice

To generate a conditional allele of Fbxw7, we employed the CRISPR-Cas9 approach to insert two FRT sites into the intron 4 and 5 of Fbxw7, respectively. Two guide RNAs targeting the integration sites (gRNA-Int5A: accgtcggcacactggtcca; gRNA-Int4A: cactcgtcactgacatcgat), two homology templates containing the FRT sequences (gRNA-Int5B: agcactgacgagtgaggcgg; gRNA-Int4B: tgcctagccttttacaagat) and the Cas9 protein were micro-injected into the fertilised mouse eggs. The offspring were screened by PCR and one line with proper integration of two FRT sites was identified.

### Analysis of public data from cancer genomics studies

Data from TCGA Research Network (Lung Squamous Cell Carcinoma (TCGA, Firehose Legacy)), including mutations, putative copy-number alterations, and mRNA Expression (mRNA expression z-scores relative to diploid samples (RNA Seq V2 RSEM; threshold 2.0), were analyzed using cBioportal software and visualized using the standard Oncoprint output (Cerami et al., 2012). The Onco Query Language (OQL) used was “USP28: MUT AMP GAIN EXP >= 2” “FBXW7: MUT HOMDEL HETLOSS EXP <= -2”. Source data was from GDAC Firehose, previously known as TCGA Provisional. The complete sample set used was (n = 178). Expression analysis was performed using GEPIA (Gene expression profiling interactive analysis) software (2017).

### Human lung tumor analysis

Human biological samples were collected, stored, and managed by the Cordoba node belonging to the Biobank of the Andalusian Health Service (Servicio Andaluz de Salud-SAS) and approved by the Ethics and Clinical Research Committee of the University Hospital Reina Sofia. All subjects gave informed consent. Pathologists assessed all samples before use. mRNA extracted from the samples was analyzed by qPCR. Primers are listed in Table 1.

**Table 1:**
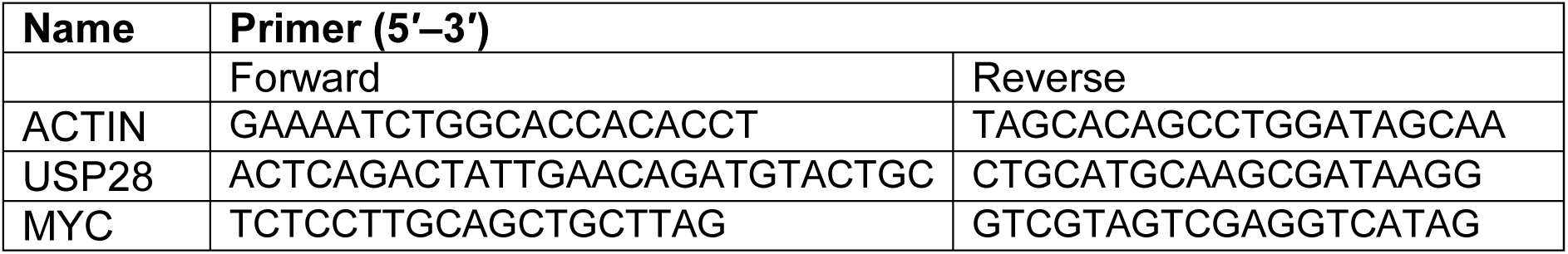
Primers for qPCR

### Tumor induction and tamoxifen treatment

Induction of NSCLC tumors was carried out in anesthetized (2-2.5% isoflurane) mice by intratracheal instillation of a single dose of 2.5×10^7^ pfu of adenoviruses encoding either the Cre recombinase (adeno-CMV-Cre) or Flp recombinase (adeno-CMV-Flp). Activation of the inducible Cre^ERT2^ recombinase was carried out by intraperitoneal injection of tamoxifen (100 μg/kg body weight) dissolved in peanut oil for 10 days.

### CT image acquisition and processing

The SkyScan-1176, a high-resolution low-dose X-ray scanner, was used for 3D computed tomography (CT). Mice were anesthetized with 2-2.5% isoflurane and CT images were acquired at a standard resolution (35 μm pixel size). The raw scan data was sorted using RespGate software, based on the position of the diaphragm, into end expiration bins. 3D reconstruction was performed using NRecon software. 3D data sets were examined using Data Viewer software.

### Mouse treatments with FT206

Nine-weeks upon Ad5-CMV-Cre infection, LSL-KRas^G12D^; Fbxw7^flox/flox^ mice were treated with FT206 (75 mg/kg) via oral gavage on day 1, 3, and 5 per week during 5 weeks. Body weights were register every week.

### In vivo pharmacology with subcutaneous graft tumors

Human LSCC tumor cell lines (NCI-H520, CALU-1 and LUDLU-1) were resuspended as single-cell suspensions at 10^7^ cells/ml in PBS:Matrigel. 100 μl (10^6^ cells total) of this suspension was injected into the flanks of immunodeficient NSG mice. When tumors were palpable, treatment with FT206 (75 mg/kg) was initiated with the same schedule on day 1, 3, and 5 per week. Tumor grafts were measured with digital callipers, and tumor volumes were determined with the following formula: (length × width^2^) × (π/6). Tumor volumes are plotted as means ± SD.

### Histopathology, Immunohistochemistry and BaseScope analysis

For histological analysis, lungs were fixed overnight in 10% neutral buffered formalin. Fixed tissues were subsequently dehydrated and embedded in paraffin, and sections (4 μm) were prepared for H&E staining or IHC. Antibodies are given in Table 2. BaseScope was performed following the manufacturer’s protocol. The *Usp28*-specific probe was custom-designed to target 436-482 of NM_175482.3; *Ppib* probe was used as a positive control (Bio-Techne Ltd).

**Table 2:** List of Reagents

| REAGENT | SOURCE | IDENTIFIER |
| --- | --- | --- |
| **Antibodies** |  |  |
| Rabbit anti-CK5 | Abcam | ab52635 |
| Rabbit anti-c-Myc | Abcam | ab32072 |
| Goat anti-GFP | Abcam | ab6673 |
| Rabbit anti-Ki67 | Abcam | ab16667 |
| Rabbit anti-TTF1 | Abcam | ab76013 |
| Rabbit anti-USP28 | Abcam | ab126604 |
| Rabbit anti-USP25 | Abcam | ab187156 |
| Rabbit anti-USP11 | Abcam | ab109232 |
| Rabbit anti-USP36 | Abcam | ab102565 |
| Rabbit anti-actin | Abcam | ab8227 |
| Rabbit anti-USP28 | Atlas | HPA006779 |
| Rabbit anti-Δp63 | Biolegend | 619001 |
| Mouse anti-c-Jun | BD Biosciences | 610326 |
| Rabbit anti-FBW7 | Bethyl | A301-721A |
| Rabbit anti-USP7 | Enzo | BML-PW0540 |
| Mouse anti-GAPDH | Invitrogen | MA5-15738 |
| Rabbit anti-Sftpc | Millipore | ab3786 |
| Rabbit anti-caspase 3 active | R&D Systems | AF835 |
| Rat anti-HA | Roche | 11666606001 |
| Mouse anti-tubulin | Sigma | T5168 |
| **Virus Strains** |  |  |
| Adeno-CMV-Cre | UI viral vector core | VVC-U of Iowa-5-HT |
| Adeno-CMV-Flp | UI viral vector core | VVC-U of Iowa-530HT |
| **Chemicals, Peptides, and Recombinant Proteins** |  |  |
| Doxycycline hyclate | Sigma | D9891 |
| Tamoxifen | Sigma | T5648 |

Tumor numbers were counted from whole lung sections: LADC and LSCC tumors were identified by Sftpc and CK5 stains, respectively. Tumor areas (μm^2^) were measured from lung sections using Zen3.0 (blue edition) software. For quantification of tumor cell death, the number of cleaved caspase-3-positive cells was counted in individual tumors per field (20x). The number of Δp63^+^, c-Myc^+^ and c-Jun^+^ cells was counted in individual tumors/10,000μm^2^. All analyses were performed uniformity across all lung sections and the whole lungs were used to derive data.

### Cell culture

Primary KF LSCC cells were cultured in N2B27 medium containing EGF (10 ng/ml; Pepro Tech) and FGF2 (20 ng/ml; Pepro Tech) (Ruiz et al., 2019). Human lung squamous cell carcinoma (NCI-H226, NCI-H520, CALU-1 and LUDLU-1) and lung adenocarcinoma (NCI-H23, NCI-H441 and NCI-H1650) lines were provided by the Francis Crick Institute Cell Services and cultured in RPMI-1640 medium supplemented with 10% FBS, 1% penicillin/streptomycin, 2mM Glutamine, 1% NEEA and 1mM Na Pyruvate. All cells were tested Mycoplasma negative and maintained at 37°C with 5% CO_2_.

### Cell treatments

Mouse KF LSCC and human LUDLU-1 cells were treated with vehicle or FT206 at different concentrations for 48hr to analyse c-Myc, c-Jun and Δp63 protein levels by western-blotting.

Primary mouse KF LSCC cells were infected with inducible-shRNAs against the Usp28 gene and then expose to Doxycycline hyclate (1µg/ml) for 48h. Cell number was counted using an automated cell counter (Thermo Fisher Scientific, Countess Automated Cell Counter).

Mouse KF LSCC and human cell lines were transfected with specific small interfering RNAs (siRNAs) against the *MYC, JUN, TP63* or *USP28* genes, using Lipofectamine RNAiMAX and 25nM of each siRNA according to the manufacturer’s instructions (Dharmacon). 72-96h later, cell number was counted using an automated cell counter. For IC50, mouse KF LSCC and human cells were treated with vehicle or FT206 at different concentrations for 72h. Cell viability was measured as the intracellular ATP content using the CellTiter-Glo Luminescent Cell Viability Assay (Promega), following the manufacturer’s instructions. IC_50_ was calculated using GraphPad Prism software.

### Western Blot Analysis

Cells were lysed in ice-cold lysis buffer (20 mM Tris HCl, pH 7.5, 5 mM MgCl_2_, 50 mM NaF, 10 mM EDTA, 0.5 M NaCl, and 1% Triton X-100) that was completed with protease, phosphatase, and kinase inhibitors. Protein extracts were separated on SDS/PAGE gel, transferred to a nitrocellulose membrane and blotted with antibodies are given in Table 2. Primary antibodies were detected against mouse or rabbit IgGs and visualized with ECL Western blot detection solution (GE Healthcare) or Odyssey infrared imaging system (LI-COR, Biosciences).

### USP28 inhibitor synthesis

Synthesis and characterization of the USP28/25 small molecule inhibitor FT206, a thienopyridine carboxamide derivative, has been described previously in the patent application WO 2017/139778 Al (Guerin, 2017) and more recent updates WO 2019/032863 (Zablocki et al., 2019) and WO 2020/033707 (Guerin et al., 2020), where FT206 is explicitly disclosed as Example 11.1.

### Cellular DUB profiling using Ub-based active site directed probes

Molecular probes based on the ubiquitin scaffold were generated and used essentially as described (Pinto-Fernandez et al., 2019; Turnbull et al., 2017). In brief, HA-tagged Ub propargyl probes were synthesised by expressing the fusion protein HA-Ub75- Intein-Chitin binding domain in E.Coli BL21 strains. Bacterial lysates were prepared and the fusion protein purified over a chitin binding column (NEB labs, UK). HA-Ub75- thioester was obtained by incubating the column material with mercaptosulfonate sodium salt (MESNa) overnight at 37°C. HA-Ub75-thioester was concentrated to a concentration of ∼1 mg/ml using 3,000 MW filters (Sartorius) and then desalted against PBS using a PD10 column (GE Healthcare). 500 μL of 1-2 mg/mL of HA-Ub75- thioester was incubated with 0.2 mmol of bromo-ethylamine at pH 8-9 for 20 minutes at ambient temperature, followed by a desalting step against phosphate buffer pH 8 as described above. Ub probe material was concentrated to ∼1mg/ml, using 3,000 MW filters (Sartorius), and kept as aliquots at -80°C until use.

### DUB profiling competition assays with cell extracts and with cells

Crude NCI-H520 cell extracts were prepared as described previously using glass-bead lysis in 50 mM Tris pH 7.4, 5 mM MgCl_2_, 0.5 mM EDTA, 250 mM sucrose, 1 mM DTT. For experiments with crude cell extracts, 50 μg of NCI-H520 cell lysate was incubated with different concentrations of USP28 inhibitor compounds (FT206 and AZ1) for one hour at 37 °C, followed by addition of 1 μg HA-UbPA and incubation for 10 minutes (Figure 4B, 4C) or 30 minutes (Figure S4A comparing FT206 and AZ1) at 37 °C. Incubation with Ub-probe was optimised to minimise replacement of non-covalent inhibitor FT206 by the covalent probe. Samples were then subsequently boiled in reducing SDS-sample buffer, separated by SDS-PAGE and analysed by Western Blotting using anti-HA (Roche, 1:2000), anti-USP28 (Abcam, 1:1000), anti-USP25 (Abcam, 1:1000), anti-GAPDH (Invitrogen, 1:1000) or beta Actin (Abcam, 1:2000) antibodies. For cell-based DUB profiling, 5×10^6^ intact cells were incubated with different concentrations of inhibitors in cultured medium for 4 hours at 37 °C, followed by glass-bead lysis, labelling with HA-UbPA probe, separation by SDS-PAGE and Western blotting as described above.

### DUB inhibitor profiling by quantitative mass spectrometry

Ub-probe pulldown experiments in presence of different concentrations of the inhibitor FT206 were performed essentially as described (Pinto-Fernandez et al., 2019; Turnbull et al., 2017) with some modifications. In brief, immune precipitated material from 500 μg-1 mg of NCI-H520 cell crude extract was subjected to in-solution trypsin digestion and desalted using C18 SepPak cartridges (Waters) based on the manufacturer’s instructions. Digested samples were analyzed by nano-UPLC-MS/MS using a Dionex Ultimate 3000 nano UPLC with EASY spray column (75 μm x 500 mm, 2 μm particle size, Thermo Scientific) with a 60 minute gradient of 0.1% formic acid in 5% DMSO to 0.1% formic acid to 35% acetonitrile in 5% DMSO at a flow rate of ∼250 nl/min (∼600 bar/40 °C column temperature). MS data was acquired with an Orbitrap Q Exactive High Field (HF) instrument in which survey scans were acquired at a resolution of 60.000 at 400 m/z and the 20 most abundant precursors were selected for CID fragmentation. From raw MS files, peak list files were generated with MSConvert (Proteowizard V3.0.5211) using the 200 most abundant peaks/spectrum. The Mascot (V2.3, Matrix Science) search engine was used for protein identification at a false discovery rate of 1%, mass deviation of 10 ppm for MS1 and 0.06 Da (Q Exactive HF) for MS2 spectra, cys carbamidylation as fixed modification, met oxidation and Gln deamidation as variable modification. Searches were performed against the UniProtKB human sequence data base (retrieved 15.10.2014). Label-free quantitation was performed using MaxQuant Software (version 1.5.3.8), and data further analysed using GraphPad Prism software (v7) and Microsoft Excel. Statistical test-s ANOVA (multiple comparison; Original FDR method of Benjamini and Hochberg) was performed using GraphPad Prism software. The MS data was submitted to PRIDE for public repository with an internal ID of px-submission #469830.

### TUBE pulldown

Endogenous poly-Ub conjugates were purified from cells using TUBE affinity reagents (LifeSensors, UM401). Cells were lysed in buffer containing 50 mM Tris-HCl pH 7.5, 0.15 M NaCl, 1mM EDTA, 1% NP-40, 10% glycerol supplemented with complete protease inhibitor cocktail, PR-619 and 1,10-phenanthroline. Lysate was cleared by centrifugation, Agarose-TUBEs were added, and pulldown was performed for 16 h at 4 °C on rotation. The beads were then washed three times with 1 ml of ice-cold TBS-T, and bound material was eluted by mixing the beads with sample buffer and heating to 95 °C for 5 min.

### Statistical analysis

Data are represented as mean ± S.E.M.. Statistical significance was calculated with the unpaired two-tailed Student’s t test, one-way or two-way analysis of variance (ANOVA) followed by multiple comparison test using GraphPad Prism software. A *P* value that was less than 0.05 was considered to be statistically significant for all data sets. Significant differences between experimental groups were: *p< 0.05, **p< 0.01 or *** p< 0.001. Biological replicates represent experiments performed on samples from separate biological preparations; technical replicates represent samples from the same biological preparation run in parallel.

## Acknowledgements

Part of this work was funded by Forma Therapeutics. This work was also supported by the Francis Crick Institute which receives its core funding from Cancer Research UK (FC001039), the UK Medical Research Council (FC001039), and the Wellcome Trust (FC001039). We thank the Discovery Proteomics Facility (led by Dr Roman Fischer) at the Target Discovery Institute (Oxford) for expert help with the analysis by mass spectrometry. Work in the B.M.K. laboratory was supported by a John Fell Fund 133/075, the Wellcome Trust (097813/Z/11/Z) and the Engineering and Physical Sciences Research Council (EP/N034295/1).

## Author Contributions

EJR, CJD, TRH, SK, DK, NJ, MW, SU, MJC, BMK and AB designed the study. EJR, LL, EMR, CDC, IR, NM and EN performed mouse genetics and in vivo experiments. EJR, TMC, AD, GV, HCS, CH and APF performed biochemical experiments with the help of APT, WWK, MG, DK and BMK. DG, JK, MK, CM, SI, JCC and CJD designed and characterised small molecule inhibitors. All authors commented on the manuscript. EJR, APF, BMK and AB wrote the manuscript.

## Supplementary figure legends

**Supplementary Figure S1, related to Figure 1.**
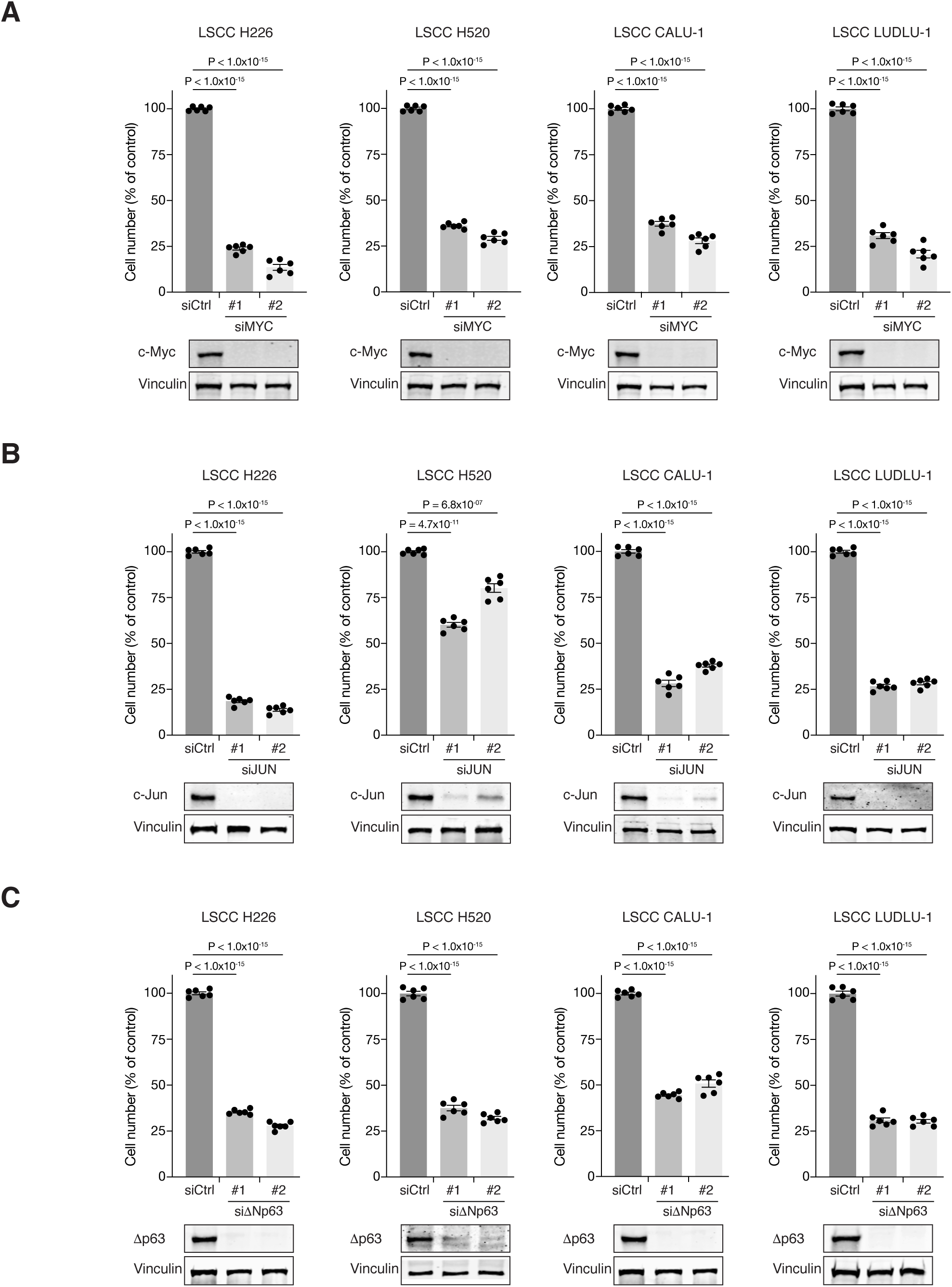
A) Graphs showing the difference in cell proliferation between control and siMYC- transfected human LSCC cell lines (NCI-H226, NCI-H520, CALU-1 and LUDLU-1). Graphs indicate mean ± S.E.M.. P values calculated using one-way ANOVA with Tukey’s multiple comparisons test. B) Graphs showing the difference in cell proliferation between control and siJUN- transfected human LSCC cell lines (NCI-H226, NCI-H520, CALU-1 and LUDLU-1). Graphs indicate mean ± S.E.M.. P values calculated using one-way ANOVA with Tukey’s multiple comparisons test. C) Graphs showing the difference in cell proliferation between control and siΔNp63- transfected human LSCC cell lines (NCI-H226, NCI-H520, CALU-1 and LUDLU-1). Graphs indicate mean ± S.E.M.. P values calculated using one-way ANOVA with Tukey’s multiple comparisons test.

**Supplementary Figure S2, related to Figure 1.**
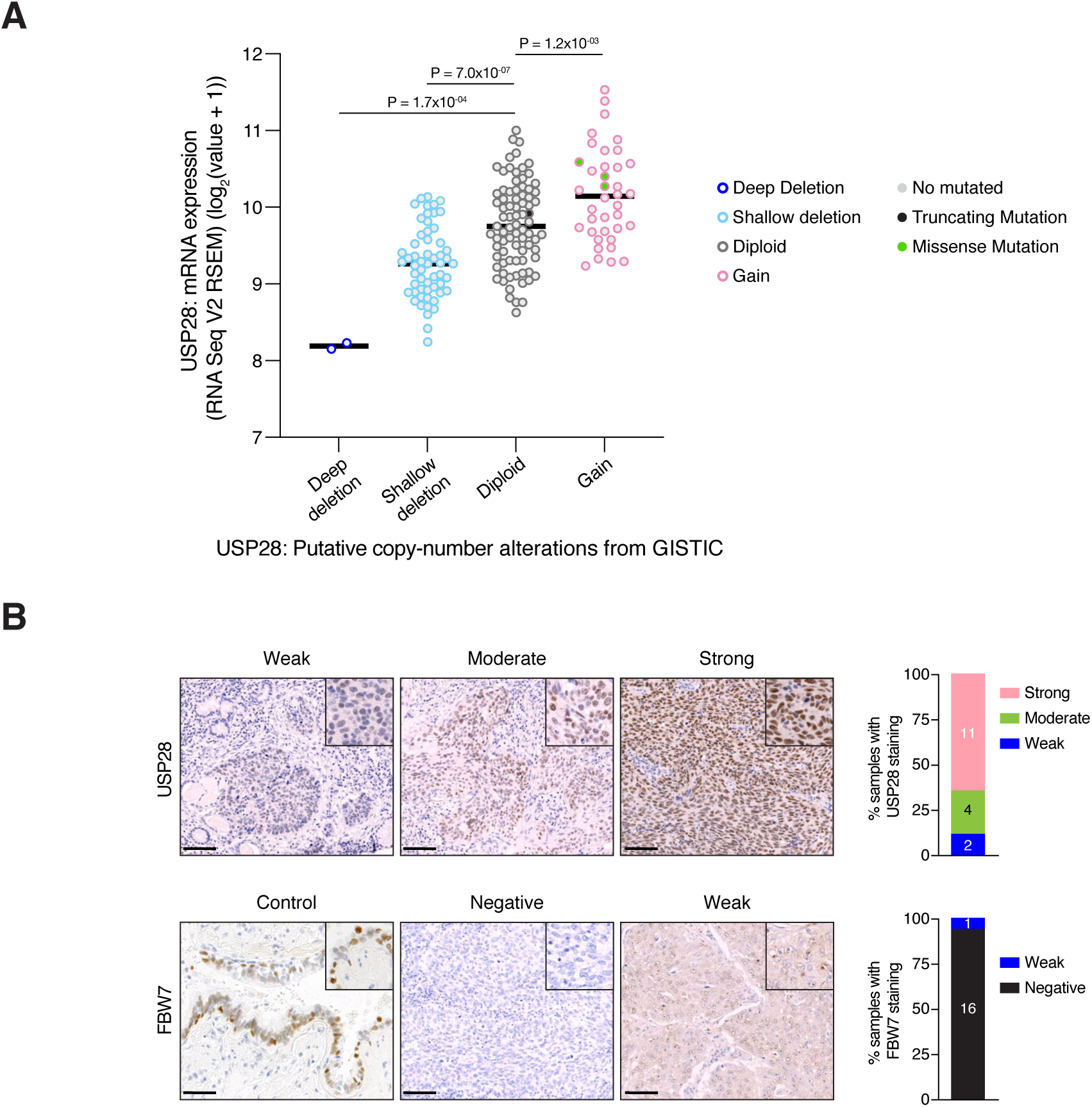
A) Dot plot showing association between the log_2_ mRNA expression (Y-axis) and copy-number alterations (X-axis) for USP28 gene. Data from TCGA were analyzed using cBioportal software. One-way ANOVA with Bonferroni’s multiple comparisons test was used to calculate P values (n = 2 Deep deletion, n = 57 Shallow deletion, n = 81 Diploid, n = 38 Gain). B) Representative human LSCC tumors stained with USP28 and FBW7 antibodies. Scale bars, 100 µm (left panel). Quantification of USP28 and FBW7 protein staining in LSCC tumors (n = 17) (right panel).

**Supplementary Figure S3, related to Figure 2.**
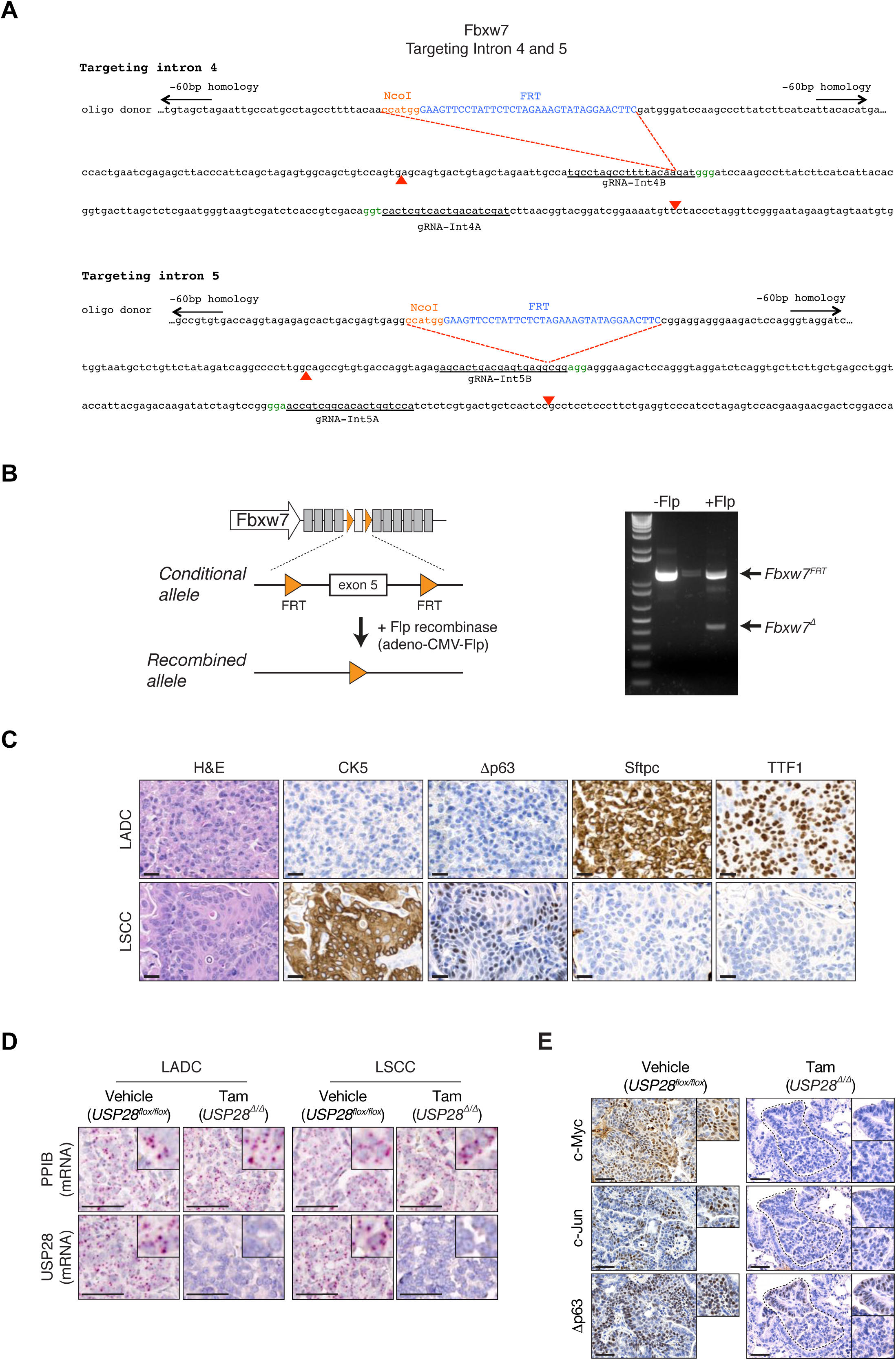
**Gene targeting strategy to generate a Fbxw7 FRT/FRT allele that can be deleted by Flp recombinase.** A) Gene targeting strategy to generate conditional Fbxw7^FRT/FRT^ animals. Two FRT sites were inserted into the intron 4 and 5 of Fbxw7 through the CRISPR-Cas9 technology. B) Schematic representation of the conditional allele (left panel). In vitro recombination assay demonstrated efficient ablation of the exon 5 upon Flp recombinase adenovirus infection (right panel). C) KFCU (FSF-<u>K</u>Ras^G12D^; <u>F</u>bxw7^FRT/FRT^; ROSA26-FSF-<u>C</u>re^ERT^; <u>U</u>sp28^flox/flox^) mice infected with adeno-CMV-Flp virus develop LADC (Sftpc^+^ and TTF1^+^) and LSCC (CK5^+^ and Δp63^+^) tumors. D) In situ hybridization of *USP28* and *PPIB* mRNA expression in vehicle- and tamoxifen-treated KFCU mice. Scale bars, 50 µm. E) KFCU tumors stained with c-Myc, c-Jun and Δp63 antibodies. KFCU mice treated with vehicle (left panel) or tamoxifen (right panel). Inserts showing c-Myc+, c-Jun+ and Δp63+ LSCC tumors in mice receiving vehicle but partial positive or negative LSCC lesions in mice receiving tamoxifen. Scale bars, 50 μm.

**Supplementary Figure S4, related to Figure 4.**
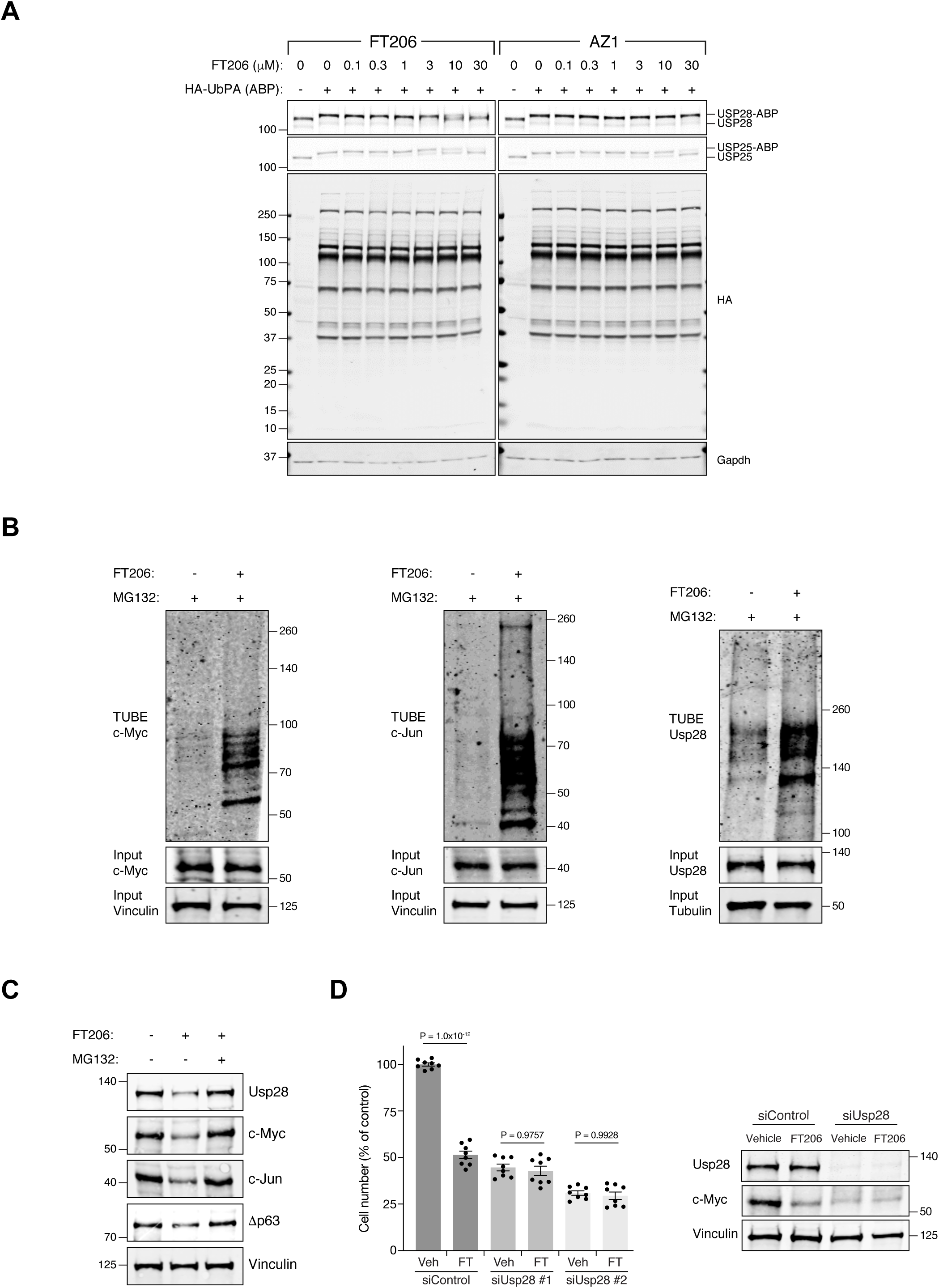
A) Comparison of USP28/25 inhibitor potency by activity-based profiling. Human LSCC H520 crude cell extracts were incubated either with AZ1 or FT206 inhibitors at indicated concentrations, followed by HA-UbPA activity-based probe (ABP) labelling. Samples were analysed by SDS-PAGE and immunoblotted using USP28, USP25, HA, and GAPDH antibodies. Inhibitor potency was reflected by competition with USP28/25-ABP adduct formation. B) TUBE pulldown of endogenous ubiquitylated c-Myc, c-Jun and USP28 in LSCC cells upon co-treatment with MG132 and FT206. C) Immunoblot of endogenous USP28, c-Jun, c-Myc and Δp63 in LSCC cells upon co-treatment with MG132 and FT206. Vinculin served as loading control. D) Graphs showing the difference in cell proliferation between control, FT206-treated and USP28-depleted LSCC cells. Graph indicates mean ± S.E.M.. One-way ANOVA with Tukey’s multiple comparisons test was used to calculate P values. Vinculin is shown as loading control.

**Supplementary Figure S5, related to Figure 5.**
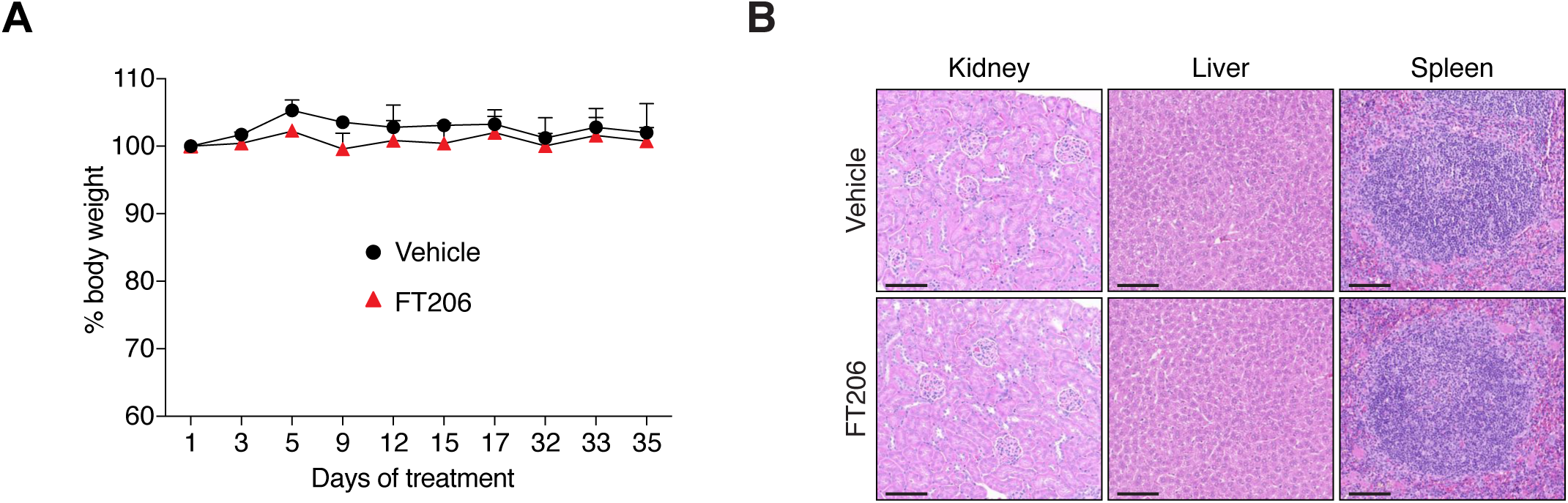
A) Monitoring tolerability in mice treated with FT206 (75mg/kg), 3 times a week for 5 weeks. Body weights of animals during the course of treatment (n = 3 vehicle, n = 3 FT206). B) Kidney, liver and spleen sections stained with H&E. Mice treated as in A. Bars, 100 μm.

**Supplementary Figure S6, related to Figure 6.**
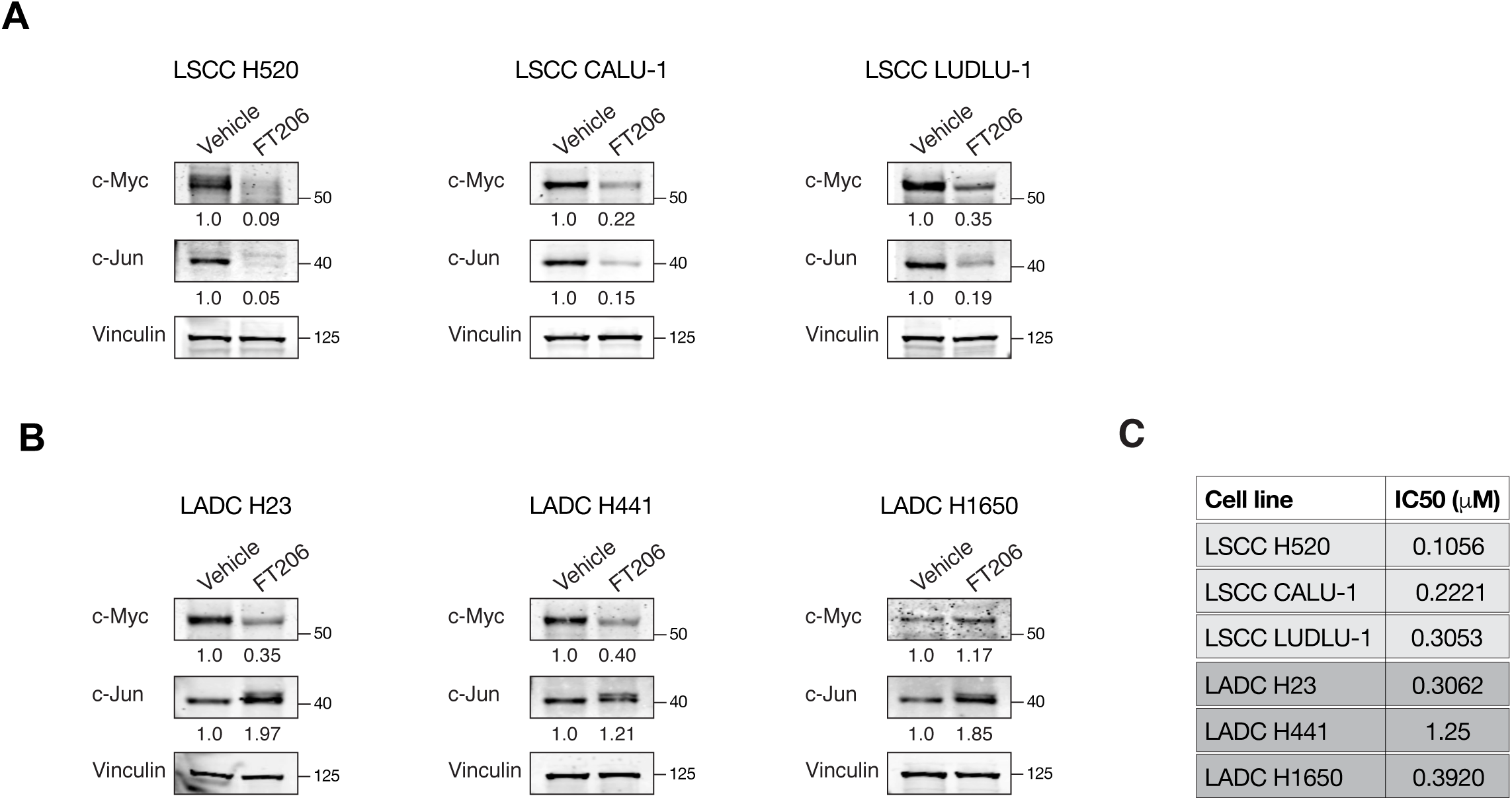
A) Immunoblot of endogenous c-Myc and c-Jun in LSCC cells upon FT206 treatment (IC50 doses display in panel C). Vinculin served as loading control. B) Immunoblot of endogenous c-Myc and c-Jun in LADC cells upon FT206 treatment (IC50 doses display in panel C). Vinculin served as loading control. C) IC50 values (doses that inhibits 50% of the cell viability) were calculated after exposure of human LADC and LSCC cells to different concentrations of FT206 compound.

